# Ventromedial prefrontal cortex is not critical for behavior change without external reinforcement

**DOI:** 10.1101/372185

**Authors:** Nadav Aridan, Gabriel Pelletier, Lesley K Fellows, Tom Schonberg

## Abstract

Cue-approach training (CAT) is a novel paradigm that has been shown to induce preference changes towards items without external reinforcements. In the task the mere association of a neutral cue and a speeded button response has been shown to induce a behavioral change lasting months. This paradigm includes several phases whereby after the training of individual items, behavior change is manifested through binary choices of items with similar initial values. Neuroimaging data have implicated the ventromedial prefrontal cortex (vmPFC) during the choice phase of this task. However, it still remains unclear what are the underlying neural mechanisms during training. Here, we sought to determine whether the ventromedial frontal cortex (VMF) is critical for the non-reinforced preference change induced by CAT. For this purpose, eleven participants with focal lesions involving the VMF and 30 healthy age-matched controls performed the CAT. We found that at the individual level, a similar proportion of VMF and healthy participants showed a preference shift following CAT. The VMF group performed similarly to the healthy age-matched control group in the ranking and training phases. As a group the healthy age-matched controls exhibited a behavior change, but the VMF participants as a group did not. We did not find an association between individual lesion patterns and performance in the task. We conclude that a fully intact VMF is not critical to induce non-externally reinforced preference change and suggest potential mechanisms for this novel type of behavioral change.

## 1. Introduction

Decision neuroscience has contributed to the understanding of maladaptive motivated behavior in conditions such as substance abuse, pathological gambling, and obesity (Bechara, 2005; Davis et al., 2004; Diekhof et al., 2008; Ernst and Paulus, 2005; Goudriaan et al., 2005). This knowledge offers opportunities for the development of new interventions to support behavior change. Most of the work in this area in humans has focused on the use of external reinforcement (Cahill and Perera, 2008; O’Doherty et al., 2004; Petry, 2010) and self-control (Baler and Volkow, 2006; Carden and Wood, 2018) to over-ride unwanted behaviors. Recently, Schonberg et al., (2014) introduced the Cue-approach training (CAT) task, in which merely associating an image with a cue and a speeded button press leads to long-lasting preference changes. The effect has been replicated in dozens of independent samples, showing changes in preferences for snack food items, unfamiliar faces, fractal art images and positive emotional images (Bakkour et al., 2016a, 2016b; Salomon et al., 2018; Zoltak et al., 2017). The CAT shows that an association of a neutral cue and a motor response with individual items (termed “Go” items) can change preference for different stimuli without external reinforcement or self-control, offering a novel avenue for addressing maladaptive choices.

The CAT procedure includes several phases. First, participants rank the stimuli to indicate their subjective preference. Based on the initial ratings, items are chosen to be associated with the button press and the cue in the following training phase. During training, the entire stimulus set is presented on the screen several times with some of the items consistently associated with the cue and the button press (“Go” items). Then, preference change is probed in a binary choice phase where two items of similar initial rankings are pitted against each other. If training did not influence preference, participants are expected to be indifferent between the two items (i.e. at chance). While the behavioral studies have shown that CAT produces a replicable group effect of about 60-65% preference of the trained Go items, the cognitive and neural mechanisms underlying this effect remain unclear. Eye-gaze data during the probe phase showed greater gaze towards Go items even when they were not chosen, compared to No-Go items. This suggests that the induced shift of preference in the CAT relies on attentional mechanisms to transform the low level visual, auditory and motor features of the training into an updated value of the associated items. Functional magnetic resonance imaging (fMRI) studies of CAT have implicated the ventromedial prefrontal cortex (vmPFC) in the probe phase of the task, showing greater activations for choices of Go items compared to choices of No-Go items modulated by the preference for individual items (Bakkour et al., 2016b; Schonberg et al., 2014). In the original study (Schonberg et al., 2014), activation of vmPFC was also observed at the end of the training phase, however, similar activation was found for both Go and No-Go items. While these studies implicate the vmPFC in the CAT during the training and choice phases, they do not reveal whether this region plays a crucial role in this preference manipulation.

Activity of vmPFC and adjacent mOFC (together termed ventromedial frontal cortex; VMF) have been implicated in representation and dynamic updating of value both in animals and humans (Wallis, 2012). In humans, activity within this area has been shown to scale with increasing subjective value across a range of stimuli types and tasks, and in some paradigms, to predict value-based choice (Bartra et al., 2013; Levy and Glimcher, 2012). Further evidence for the critical role of VMF in value-based decisions comes from lesion studies. Participants with VMF damage show impaired performance in flexible value learning tasks (Fellows and Farah, 2003; Tsuchida et al., 2010) and make less consistent preference judgments compared to healthy controls (N. Camille et al., 2011; Fellows and Farah, 2007; Henri-Bhargava et al., 2012). Importantly, VMF damage was recently found to disrupt biasing of attention to rewarding features of the environment, suggesting that this area is critical to the interplay of attention and value in decision-making (Vaidya and Fellows, 2015).

Activation within VMF during choice in the fMRI studies of CAT suggests that this region might be necessary for the value update that underpins preference change in CAT. Alternatively, other structures (e.g. the visuomotor network) encode the value update and VMF is merely active during the choice of the preferred item, reflecting the updated value rather than making a causal contribution to the preference change. These two models propose different roles for VMF in CAT (Fig. 1); in the first, this region dynamically assigns credit following low-level attentional training. In the second, VMF is not involved in modifying value, but is involved in value representation during choice. These models make different predictions regarding the effects of VMF damage on CAT performance: if intact VMF is necessary for the CAT effect, individuals with VMF damage will show an attenuated or absent shift of preference following CAT. Alternatively, if VMF is not necessary for the value updating during training or value retrieval during choice, VMF damage will not affect preference shifts following CAT. In the current study, we tested these competing hypotheses by examining whether focal VMF damage affects the shift of preferences observed following CAT. Understanding the role VMF plays in behavior change with CAT will shed light on the one hand on this novel non-externally reinforced procedure and on the other on the role VMF plays in value construction and assignment during value based-decision making more generally.

**Figure 1.**
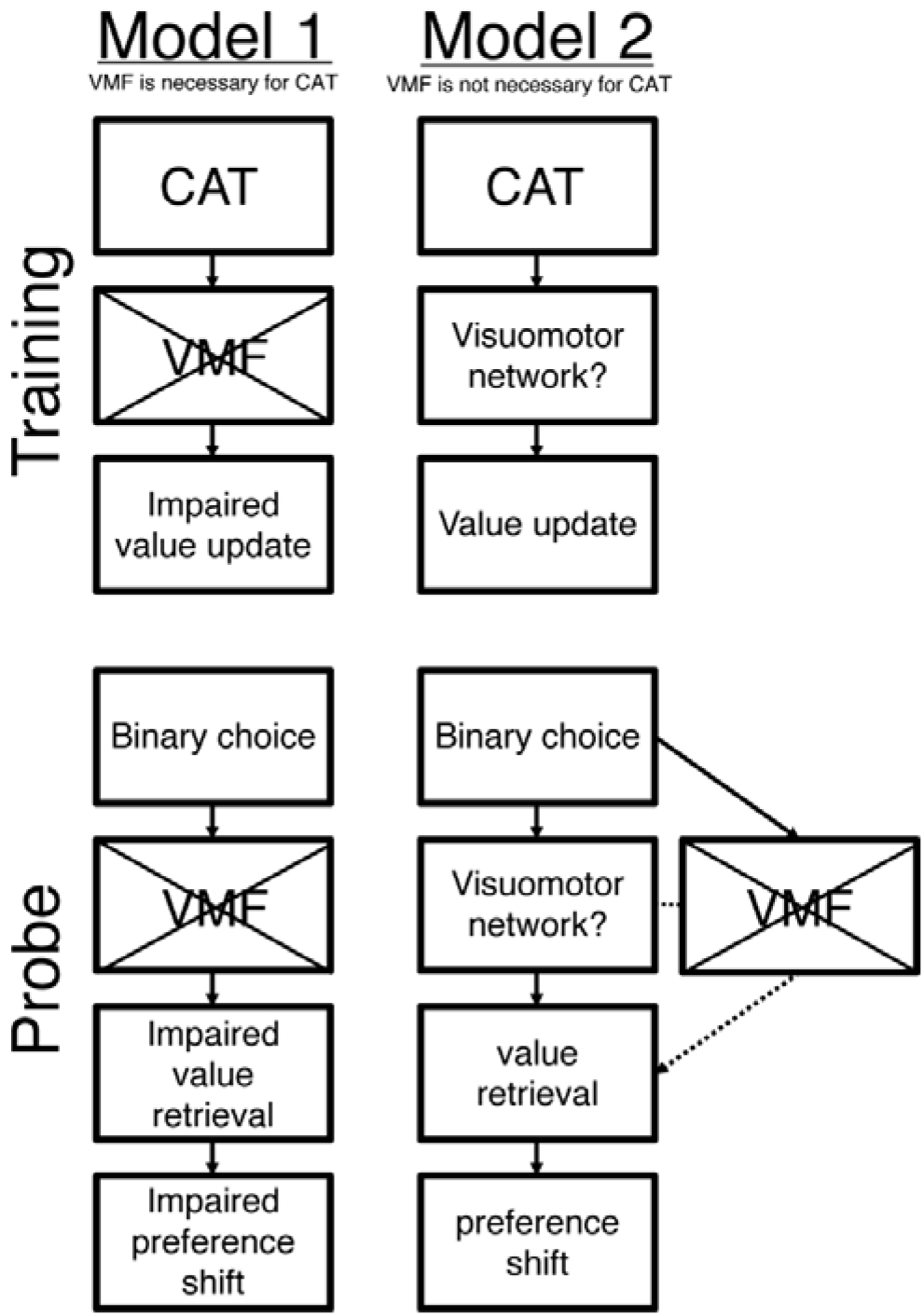
Putative models for the mechanisms underlying CAT.

## 2. Materials and methods

### 2.1. Participants

Participants with focal lesions involving the orbitofrontal cortex (OFC) and ventromedial prefrontal cortex (vmPFC), together referred to here as ventromedial frontal: VMF (*N* =11, mean age = 59.4 [39-78] years, 5 males), were recruited from the Cognitive Neuroscience Research Registry at McGill University. All had fixed, circumscribed lesions of at least 6- months duration (mean duration = 9.3 [5.4-16.5] years). Lesions were due to ischemic stroke, tumor resection, or aneurysm rupture. Thirty age-matched healthy control participants were recruited through local advertisements in Montréal. They were free of neurological or psychiatric disease and were not taking any psychoactive drugs. One control participant was excluded from the analysis due to extremely inconsistent choices (choice prediction accuracy = 0.52 z = −3.62; see Results for details). For the 29 included in this group, the mean age was 60.5 [44-79] y and 15 were females. All participants provided written, informed consent in accordance with the Declaration of Helsinki and were paid a nominal fee for their time. The *Missing data from 1 subject. The study protocol was approved by the local Research Ethics Board.

**Table 1.** VMF group neuropsychological screening test performance [mean (SD)]

|  | **Incidental**<br>**memory**<br>(accuracy) | **Fluency –**<br>**animals**<br>(words/1 min) | **Fluency –**<br>**F**<br>(words/1 min) | **Backwards**<br>**digit span** | **Sentence**<br>**comprehension**<br>(accuracy) |
| --- | --- | --- | --- | --- | --- |
| **VMF** $N = 11$ | 0.81 (0.14)* | 17.6 (2.4) | 10.6 (5.6) | 2.8 (0.9) | 0.99 (0.02)* |

\*Missing data from 1 subject.

### 2.2. Lesion Analysis

Individual lesions were traced from the most recent clinical computed tomography or magnetic resonance imaging onto the standard Montreal Neurological Institute (MNI) brain using MRIcro software (Rorden & Brett, 2000; www.mccauslandcenter.sc.edu/mricro/) by a neurologist experienced in imaging analysis and blind to task performance. MRIcron (www.nitrc.org/projects/mricron) was used to generate lesion overlap images (Fig 2).

**Figure 2.**
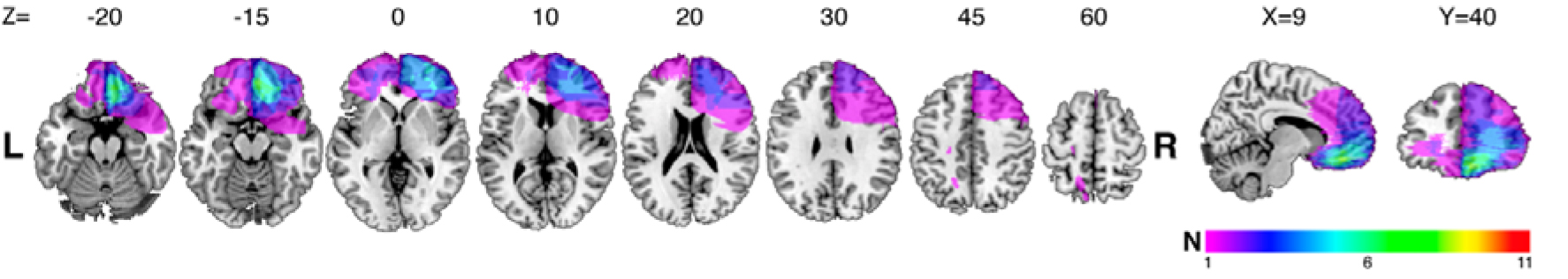
Lesion extent and overlap. Lesion overlap across the VMF participants is indicated by color bar, MNI brain slice coordinates are indicated by XYZ.

### 2.3. Procedure

Sixty identically-sized color images of computer-generated fractal art images served as the stimuli (“Fantastic Fractals,” 2013). The experiment was run using MATLAB (Mathworks, Inc. Natick, MA, USA) on a 21-inch screen.

#### 2.3.1. Binary ranking

A forced-choice binary ranking procedure was used to estimate participants’ baseline subjective preferences for each of the stimuli. In this task, 60 stimuli were randomly paired to form 300 unique pairs. For each pair of stimuli, participants had 2500 ms to choose their preferred stimulus, followed by a 500 ms choice confirmation screen and 500 ms fixation cross (Fig. 3A). Based on the assumption of choice transitivity from rational choice theory (Regenwetter et al., 2011; Von Neumann and Morgenstern, 1944), we used the outcomes from the set of binary choices to infer individual preferences for the presented set of stimuli. That is, if stimulus A is preferred over B and stimulus B is preferred over C, then their respective ranks are A>B>C. We used the Colley Matrix algorithm (Colley, 2002), designed to solve ranking problems with limited number of binary outcomes to maximize ranking validity and specificity. This procedure resulted in a ranked list of the 60 stimuli, based on each participant’s individual preferences. Colley Matrix ranking scores typically range from 0 (least liked) to 1 (most liked), with a fixed mean of 0.5. An intransitive choice pattern is characterized by densely distributed scores around the center of 0.5, while a distinct preference pattern leads to more distributed ranking scores. From these rankings, we quantified a transitivity score for each participant as the standard deviation of the participant’s ranking scores.

**Figure 3.**
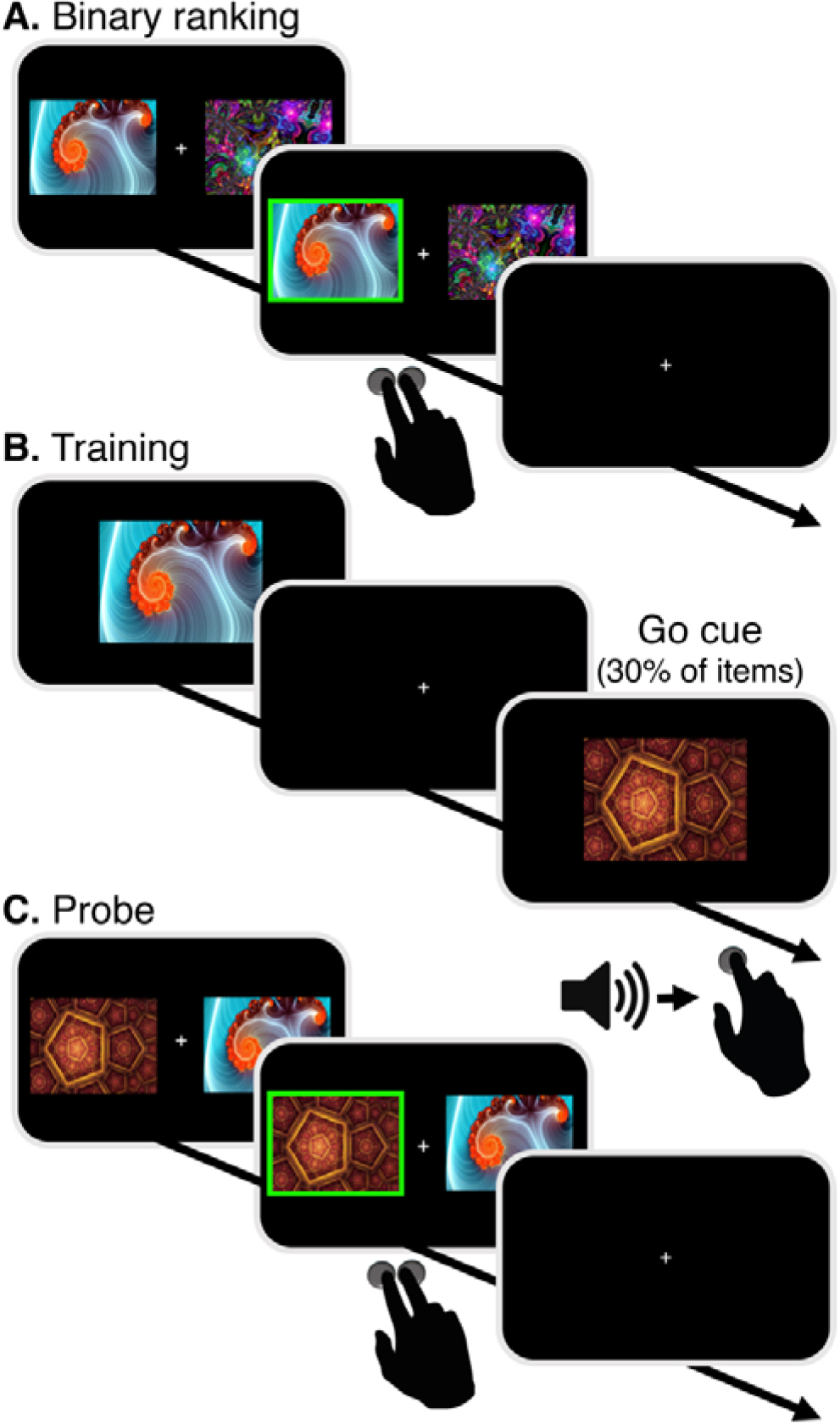
Procedure. A) Binary choices between pairs of 60 fractal images were used to obtain rankings; B) Training of Go-items, consistently paired speeded button presses cued by a neutral tone across 16 runs; C) Probe: binary choices between pairs of Go and No-Go items with similar initial ranking.

#### 2.3.2. Cue-approach training

Following the baseline evaluation of subjective preferences using the binary ranking procedure, participants underwent 16 training runs of the cue-approach training procedure. Twelve of the 60 stimuli (6 high-value, mean rank = 12.5 and 6 low-value mean rank =48.5) were consistently associated with a Go auditory-cue (Fig. 3B). Each stimulus in the training set was presented individually on the screen for 1000 ms, once during each training run. The cue appeared following an adaptive delay according to participant performance, as in previous CAT studies (Schonberg et al., 2014), this ensured maintaining a similar difficulty level throughout the training phase and across individuals. Stimuli were randomly ordered and followed by a jittered fixation cross with an average duration of 2000-ms (range of 1000- 6000-ms, 1000- ms intervals).

#### 2.3.3. Probe

Preference change following CAT was evaluated in a probe phase. On each probe trial, two items appeared to the right and left of a central fixation cross and participants were asked to select their preferred stimulus. In each pair, both items were of similar initial value (either high-value or low-value), but only one item was a Go item, i.e. associated with a cue during training. For each pair, participants had 1500-ms to select their preferred stimulus, followed by a 500-ms choice confirmation and a fixation cross for a jittered duration with an average of 3000-ms (range of 1000-11000-ms, 1000-ms intervals; Fig. 3C). In addition to these comparisons, as in previous CAT experiments, ‘sanity check’ trials were also incorporated in the probe phase to measure preferences consistency. In the ‘sanity check’ trials, participants were asked to choose between pairs of items in which one item was of initial high-value and the other of initial low-value (both Go or both No-Go items), to validate the stability across time of the initial preference evaluation. The probe phase included two runs with 152 total trials, with all unique probe pairs presented in a random order in each run.

#### 2.3.4. Memory

At the end of the experiment, participants performed two sequential memory tasks. The first assessed memory for fractals presented during the experiment compared to novel items (Old/New). The second assessed whether participants remembered which images were associated with the cue (Go/No-Go).

### 2.4. Statistical analyses

Binary choice outcomes were analyzed using a mixed-model logistic-regression (R package lme4 v1.1-13). Group means were compared by a two-sample permutation test (R package Deducer v0.7-9), and 95% confidence intervals were estimated using data bootstrapping.

### 2.5. Data sharing

Behavioral data and analysis codes are available at osf.io/d8ceg/.

## 3. Results

See Table 2 for summary of behavioral results for all tasks.

### 3.1. Binary ranking

For each participant, we estimated choice consistency as the prediction accuracy of a “leave one out” model. For each of the 300 binary choices, we applied the Colley-matrix algorithm to the remaining 299 choices, which yielded the ranks of the 2 competing items in this trial. For each participant, we defined the choice consistency index as the proportion of trials in which the highest ranked item was chosen (% correct choice predictions). We did not find a difference in choice consistency between the groups (t_(16.9)_ = −0.08, p = 0.96, 95% CI [-4.5, 4.1]; Fig. 4A). We found a difference between the control and VMF groups in the mean reaction time during binary choices such that overall, the VMF group took slightly longer to make choices (t = −2.37, p = 0.026, 95% CI [-239, −25]; Fig. 4B). Examining the difference in RT of consistent and inconsistent choices, we did not find a difference between the control and VMF groups (t_(30.1)_ = 0.70, p = 0.47, 95% CI [-45, 96]).

**Figure 4.**
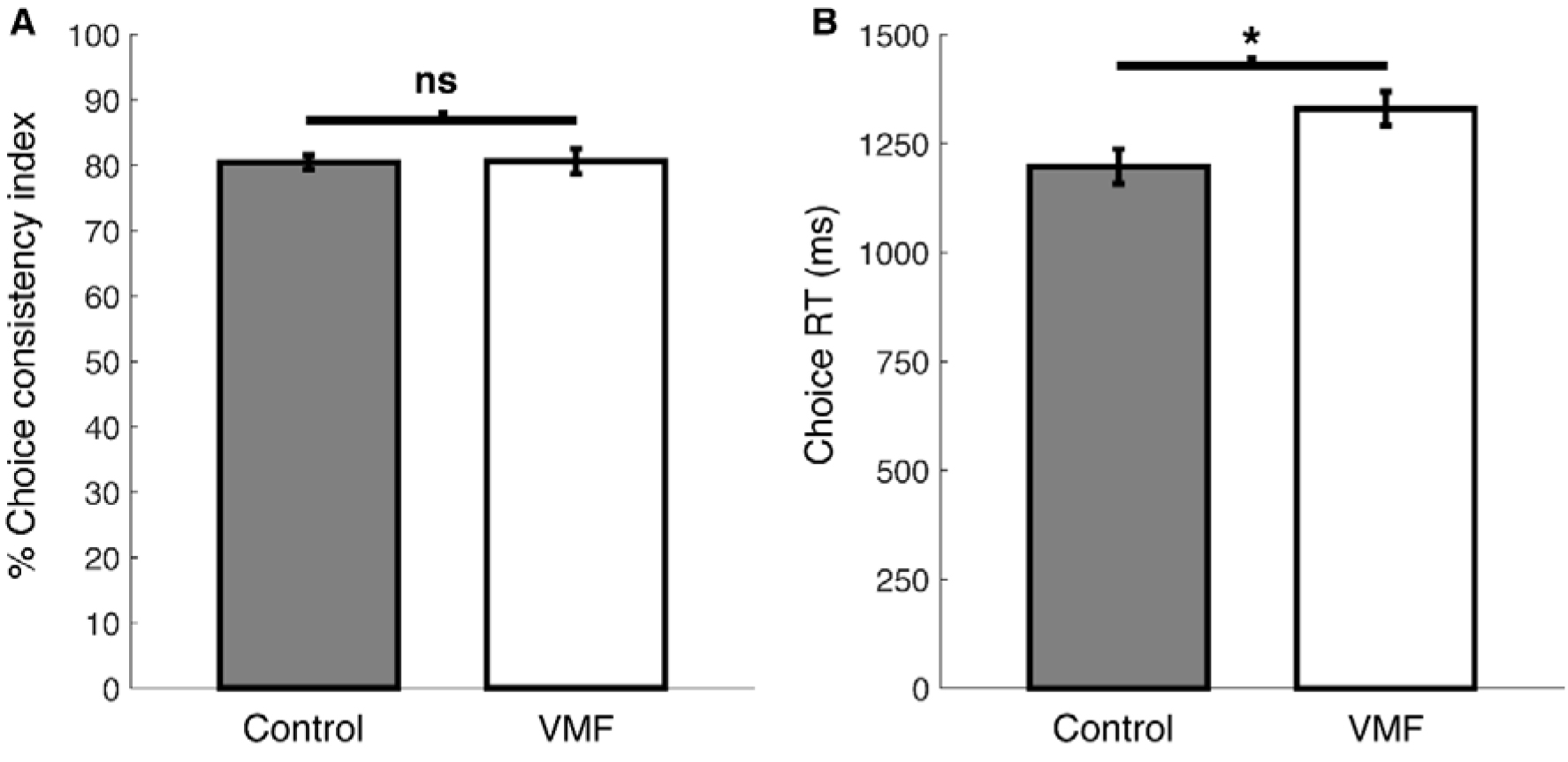
Binary ranking. A) There was no difference in choice consistency between the control and VMF groups B) The VMF group took longer time to make choices. Error bars represent standard error of the mean.

We next examined the relationship between choice-difficulty (items’ rank difference) and reaction time. In both groups, there was a negative correlation between the rank difference of a given pair of items and choice reaction time, such that the larger the rank difference between the two items, the faster the choice was made. This correlation was similar in control and VMF groups (t_(21.6)_ = 1.26, p = 0.29, 95% CI [-0.05, 0.09]).

**Table 2.**
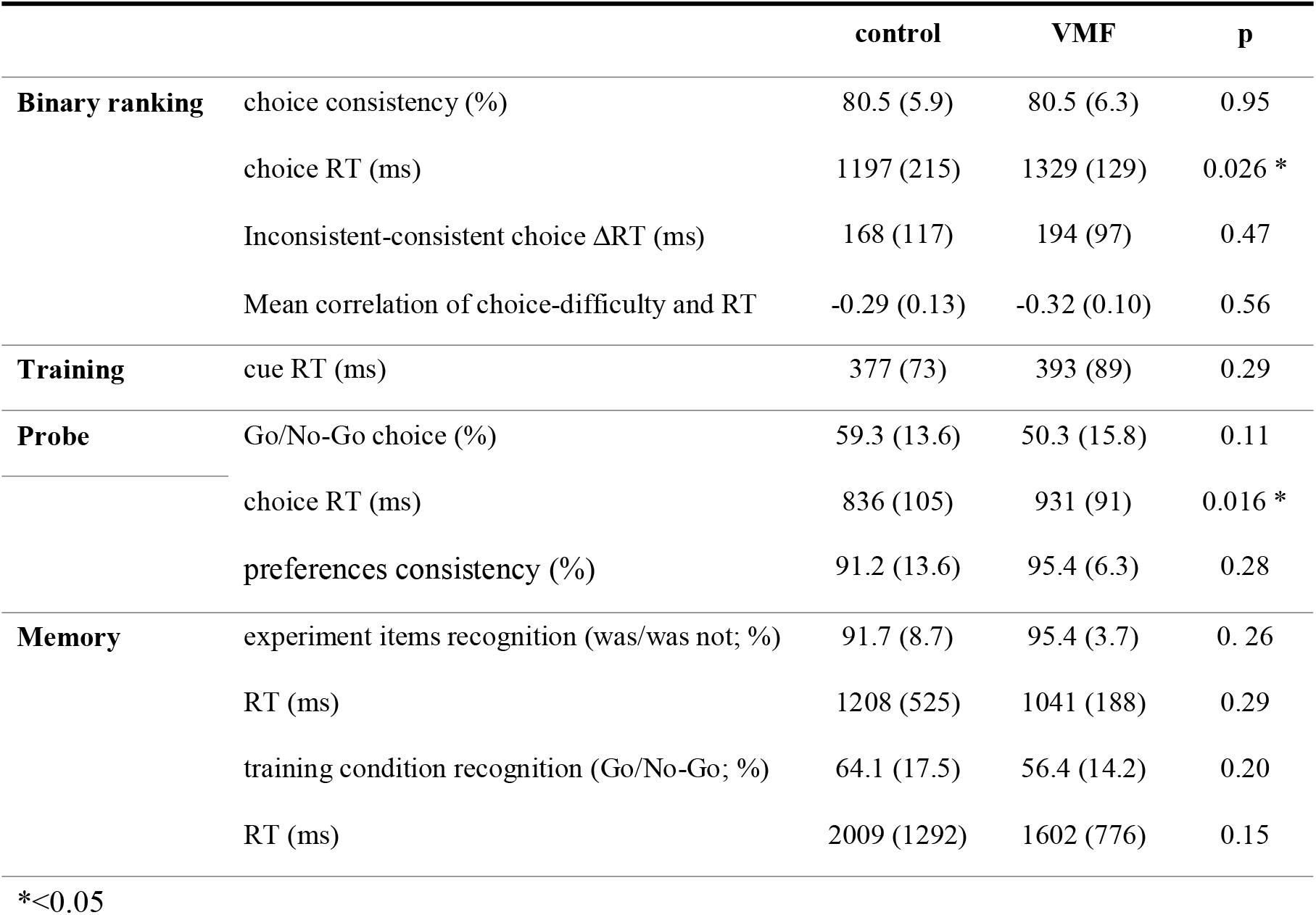
Behavioral results [mean (SD)].

|  |  | control | VMF | p |
| --- | --- | --- | --- | --- |
| **Binary ranking** | choice consistency (%) | 80.5 (5.9) | 80.5 (6.3) | 0.95 |
|  | choice RT (ms) | 1197 (215) | 1329 (129) | 0.026 * |
| | Inconsistent-consistent choice ΔRT (ms) | 168 (117) | 194 (97) | 0.47 |
|  | Mean correlation of choice-difficulty and RT | -0.29 (0.13) | -0.32 (0.10) | 0.56 |
| **Training** | cue RT (ms) | 377 (73) | 393 (89) | 0.29 |
| **Probe** | Go/No-Go choice (%) | 59.3 (13.6) | 50.3 (15.8) | 0.11 |
|  | choice RT (ms) | 836 (105) | 931 (91) | 0.016 * |
|  | preferences consistency (%) | 91.2 (13.6) | 95.4 (6.3) | 0.28 |
| **Memory** | experiment items recognition (was/was not; %) | 91.7 (8.7) | 95.4 (3.7) | 0.26 |
|  | RT (ms) | 1208 (525) | 1041 (188) | 0.29 |
|  | training condition recognition (Go/No-Go; %) | 64.1 (17.5) | 56.4 (14.2) | 0.20 |
|  | RT (ms) | 2009 (1292) | 1602 (776) | 0.15 |

\*<0.05

### 3.2. Cue approach training

There were no differences between the groups in the button-press reaction time to the tone cue (calculated as time after cue) during training between the groups (t_(15.3)_ = 1.26, p = 0.29, 95% CI [-73, 41]). *<0.05

### 3.3. Probe

#### 3.3.1. Group analysis

To assess preference changes following training, we analyzed the proportion of probe trials in which participants preferred the Go items over the No-Go items, using a two tailed repeated measures logistic regression. In each pair, both items were of similar initial preference based on the baseline evaluation phase. As in previous studies, we hypothesized that the cue approach effect would enhance preferences for the Go items above the chance level of 50% of trials (log-odds = 0; odds-ratio = 1).

Following cue-approach training, control participants consistently preferred the Go items over the No-Go items during probe (mean proportion = 59.28% (13.6), OR = 1.58, 95% CI [1.17, 2.16], p = 0.002; Fig. 5A). In the VMF group, participants did not show a preference for Go over No-Go items during probe (mean proportion = 50.39% (15.8), OR = 1.03, 95% CI [0.664, 1.59], p = 0.896; Fig. 5A, see Fig. 5B for individual VMF participants lesion maps and color codes). There difference between the control and VMF groups in Go vs. No-Go choice proportion was not significant (∆_(control-VMF)_ mean proportion = 8.89%, OR = 0.652, 95% CI [0.378, 1.12], p = 0.11).

**Figure 5.**
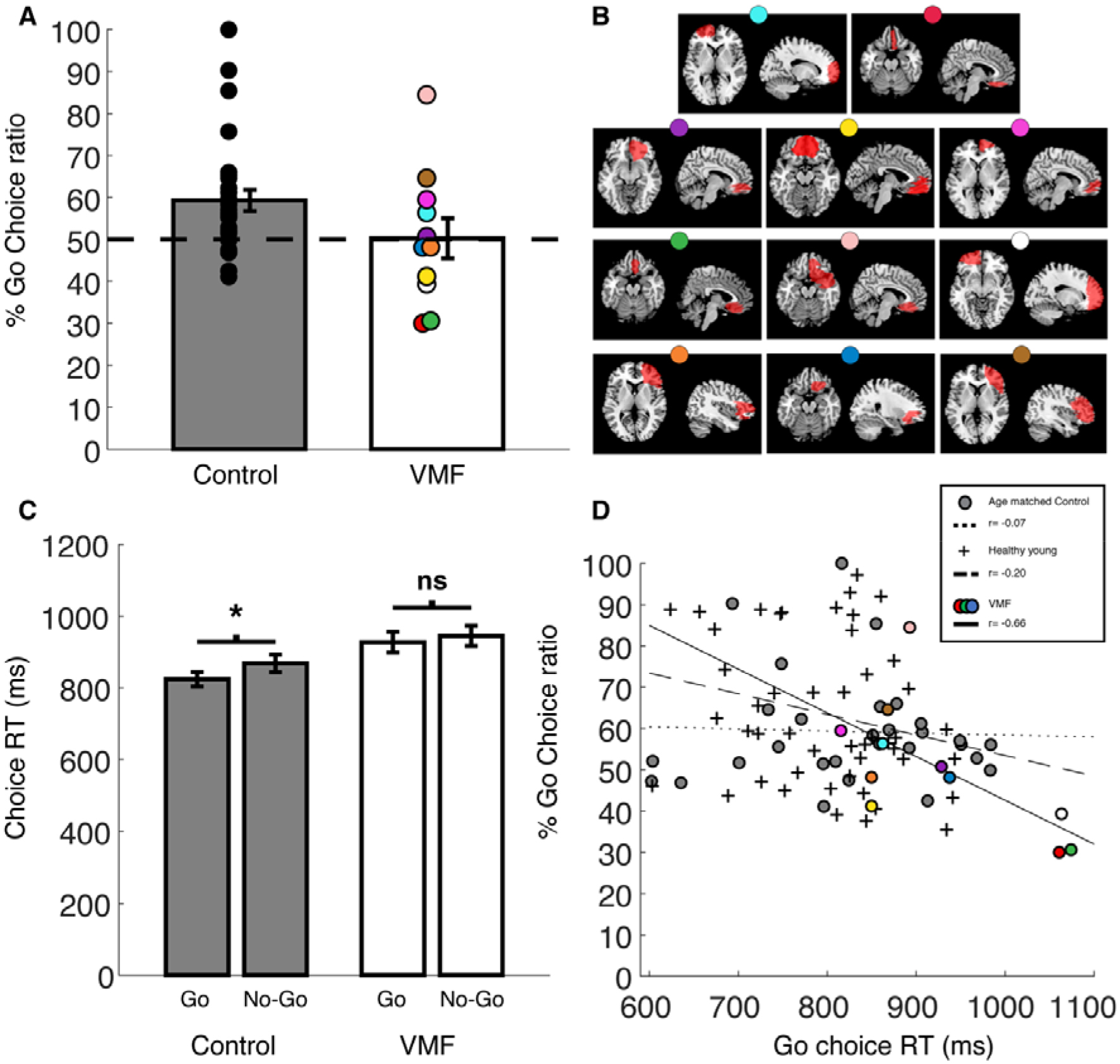
Probe. A) Ratio of Go to No-Go choices. B) Color coded lesion maps of individual VMF participants. C) Probe choice-time time. D) Correlation across participants between Go choice-time and ratio of Go to No-Go choices.

### 3.3.2. Individual participant analysis

For each participant, we calculated the individual probability of obtaining a preference shift. We defined the threshold of individual learning based on the binomial distribution compared to chance: 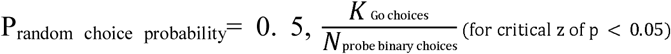 = Go choice-ratio ≥ 57.64%. This resulted in three of the 11 VMF participants showing Go choice-ratio larger than the critical Go choice-ratio and 12 participants out of 29 in the control group. Four VMF participants were below chance, compared to one in the control group (Go choice-ratio ≤ 42.36%).

### 3.3.3. Reaction time during probe

We found a difference between the control group and VMF group in choice reaction time during the probe phase with slightly longer RTs for the VMF group (t_(20.7)_ = −2.79, p = 0.011, 95% CI [-160, −32]). In the control group the choice time of the Go items was faster than choice time of the No-Go items (t_(28)_ = −2.88, p = 0.008, 95%, CI [-75, −13]). In the VMF group there was no difference in choice time of the Go and No-Go items (t_(10)_ = −1.28, p = 0.23, CI [-47, −13]; Fig. 5C). Choice reaction time and preference are known to be linked (Mir et al., 2011; Shadlen and Shohamy, 2016). Considering the difference between the control and VMF groups in choice reaction time in the probe, we explored the relation between choice reaction-time of Go items and the degree of preference shift toward the cued Go items across participants. In the control group there was no correlation (N = 29, r = −0.04, p = 0.84). However, in the VMF group, we found a significant negative correlation (N = 11, r = −0.64, p = 0.03) such that the faster the participant chose the Go items the greater the ratio of Go over No-Go items chosen (Fig. 5D).

### 3.3.4. Probe preference consistency

There were no differences between the groups in preference consistency (ratio of choosing high value items to low value items) during probe between the groups (t_(36.2)_ = −1.35, p= 0.28, 95% CI [-0.10, 0.02]).

### 3.4. Memory

At the end of the experiment, we tested participants’ memory of the items (were the items presented in the experiment, or novel) and of the training condition for the items (were the items associated with a cue and button-press response, or not). We did not find a difference in the proportion of items recognized between the control and VMF groups (t_(37.3)_ = −1.48, 95%, p = 0.28, CI [-8.7, 3.7]). We did not find a difference in the recognition ratio of the experiment items’ training condition between groups (t_(15.3)_ = 3.92, p = 0.29, 95% CI [-3.3, 18.5]). There was no difference between the control and VMF in the reaction time to recognition of the experiment items (t_(38)_ = 1.47, p = 0.29, 95% CI [-52, 384]), and no difference between the control and VMF in the reaction time to recognition of the experiment items’ training condition (t_(22.8)_ = 1.15, p = 0.38, 95% CI [-251, 1064]).

## 4. Discussion

In this study, we examined whether VMF is critical for behavior change that does not rely on external reinforcement with the novel cue approach task. We aimed to differentiate between two potential underlying models of the effect: in one, VMF has a crucial role in the transformation of the visual, auditory and motor features of training into an enhanced value of the cued items. In the other, preference change with CAT does not critically depend on VMF. We tested these competing models by studying participants with VMF damage and healthy age-matched control and examined performance across the different phases of the CAT task. We found that the same proportion of participants in the VMF group and healthy aged-matched controls showed an individual preference change effect.

As a group, the VMF participants performed similarly to the healthy aged-matched control in all task phases. The VMF participants in our study were able to make consistent preference-based choices between abstract fractal images in the binary ranking phase, similar to the control group. In the training phase, the VMF participants were indistinguishable from controls, showing intact ability to rapidly respond with a button press to the auditory cue. Thus far the CAT was tested at the group level (e.g. Schonberg et al.,2014, Salomon et al., 2018). Here, we replicated this effect for the first time in elderly controls. The VMF participants did not differ significantly from the controls, but did not show the effect as a group. This was due to a subset of VMF participants that showed the opposite effect (preference shift towards No-Go items). Thus, although inference is limited by the variability of the VMF participants, the presence of the effect in a similar proportion of individuals with VMF damage and healthy controls argues against a critical role for this region in this form of non-externally reinforced preference change.

The findings in healthy older controls confirm prior work in younger participants: the CAT effect is reliably evident at the group level (Bakkour et al., 2016a, 2016b; Schonberg et al., 2014; Veling et al., 2017; Zoltak et al., 2017), but not all individuals show the effect. An individual level analysis found that a similar proportion of the VMF participants and control group participants showed the expected CAT effect. In addition, four VMF participants showed an opposite individual effect (i.e. greater than chance preference for the No-Go items), a pattern seen in only 1/29 controls. Averaging across these two patterns in VMF participants yielded an absence of the group effect.

The lack of differences between the groups and the presence of an effect for at least some of the VMF participants support the suggestion that VMF is not critical to induce the effect, in line with previous imaging findings that showed a vmPFC response to both Go and NoGo items at the end of training (Figure 4 in Schonberg et al., 2014). In addition, the ventromedial frontal cortex encompasses several cortical areas (Rudebeck et al., 2008). It is possible that only damage to a specific sub-structure within it, is related to CAT effect. Unfortunately, a careful analysis of the individual lesions compared to the individual behavioral effects (see Figure 5) did not provide evidence for that possibility.

The VMF participants in our study could make consistent preference-based choices between abstract fractal images, similar to the control group. This finding supports previous reports of consistent value-based ratings and choices of artwork stimuli in individuals with VMF damage over short time frames (Vaidya and Fellows, 2015). In contrast, several studies reported that individuals with VMF damage were less consistent in choices of various stimuli including food items, images of human, animals and landscapes and colors (N. Camille et al., 2011; Fellows and Farah, 2007; Henri-Bhargava et al., 2012) and impaired in transitive (Koscik and Tranel, 2012) and associative inference-learning (Spalding et al., 2018). In our study, we found no differences between the VMF and control groups in choice consistency during the binary ranking phase and over the course of the 1-hour experimental session. Together with the wider literature, this suggests that VMF is required for consistent valuation of only certain categories of stimuli. The mechanisms underlying this observation remain to be clarified (Vaidya et al 2017).

One of the limitations of the current study is that we did not pre-register the sample size of the VMF group and thus we could be under powered. However, the sample size in our study was similar in size to other VMF studies that showed impairments in reinforcement learning, preference judgments and emotional processing (Nathalie Camille et al., 2011; Fellows et al., 2005; Gillihan et al., 2011; Spalding et al., 2018; Vaidya et al., 2017). Furthermore, the results obtained in both groups suggest that some individuals show the non-reinforced change while others do not. Thus, even with the current sample size we believe we can conclude that there are VMF lesion participants that show an entirely intact effect and thus this region is not critical to induce the effect. As the range of choices showing preference for Go items in this task is relatively narrow (above 58% choice) then it is hard to estimate if we obtained a blunted dilution effect.

Interestingly, we found that the VMF participants were slower to make choices (regardless of choice difficulty or choice consistency with overall individual preference) both in the binary ranking phase and in the probe phase compared to the control group. There are conflicting reports regarding choice reaction-times and VMF damage. Rogers et al., (1999) found longer deliberation time in participants with damage to OFC compared to participants with damage to dorsal or lateral prefrontal-cortex and healthy control in a decision-task that involved choices between two stimuli groups that represented varying win probabilities. Other studies reported choice reaction time was unaffected by VMF damage (Fellows, 2006; Fellows and Farah, 2007). In a previous study, it has been shown that allowing participants extra time to make a choice eliminated the CAT effect (Veling et al., 2017). In the control group of our study, choices of Go items were faster than choices of No-Go items whereas in the VMF group there was no such difference. When we explored whether choice time of Go items could explain the variability of the Go to No-Go choice ratio, we indeed found a correlation only in the VMF group. Thus, it is possible that longer choice times in the VMF group led to the lack of group effect. It is unclear whether the longer reaction time also led 4 VMF participants to show the opposite choice effect.

Beyond our main aim to shed light on the neural mechanism of CAT, this study is novel in several additional aspects. First, we show for the first time that a sample of healthy elderly participants (mean age 60.5) express a CAT effect (at the group level), similar to that observed in previous samples of young healthy adults (mean age 24; Salomon *et al.*, 2018). Second, we provide further support to previous findings that people with VMF damage can make consistent preference judgements, at least under specific conditions.

Here we adopted the triangulation approach to study the neural basis of a novel non-reinforced behavioral change task (Munafò and Davey Smith, 2018). Although we did not obtain clear cut findings, we conclude that there are no strong evidence that intact VMF is critical to induce non-externally reinforced preference change, by the mere association of cues and button presses. Between-participant variability in the CAT effect, seen in both healthy and VMF participants, limits the strength of this conclusion. The fact that the VMF group performed similarly in all other components of the procedure speaks to the ability of this group to perform value-based choices and valuation, at least of fractal art images. The finding of a correlation between reaction times and the degree of effect in the task calls for a better understanding of individual differences that underlie non-reinforced behavioral change with CAT. Further research is needed to fully determine the underlying neural mechanisms of the CAT and the role value representation in the VMF is playing within it toward future potential applications of this task.

## Acknowledgements

This work was supported by the Israeli Science Foundation (ISF number 1798/15 and 2004/15) and the European Research Council (ERC) under the European Union’s Horizon 2020 research and innovation program (grant agreement n° 715016) granted to Tom Schonberg. Nadav Aridan was supported by the Eldee Foundation and the Bloomfield family of Montréal, Canada as part of the Brain@McGill and Tel Aviv University Collaboration.

